# *In-Silico* analysis of Human Insulin Gene for Mirror repeat sequences

**DOI:** 10.1101/2023.07.10.548352

**Authors:** Manisha Yadav, Barkha Sehrawat, Pooja Yadav, Jyoti Kumari, Priyanka Yadav, Shivani Yadav, Rachna Yadav, Amrita Singh, Sandeep Yadav

## Abstract

The genetic material contains all the instructions necessary for the development, functioning & regulation of biological processes. This is also considering as a regulatory molecule. The structure constitute by its unique pattern of bases make it unique for every living being consider from bacteria to human. The varied pattern of bases in genomes or genes represents different types of repeats. These repeats types constitute the major fraction of the genomes. Among varied types Mirror repeats (MR’s) play crucial roles in the genome especially in formation of H-DNA structure by Non Watson pairing. Some studies suggest that mirror sequences responsible for neurological diseases also. Now days in modern era of technology *in silico* techniques were develop to extract any type of repeat sequences from any gene or genome. The major focus of current work is on identification of mirror repeats (MR’s) from human insulin gene sequence using an *in silico* approach. The approach is refer to as FPCB (FASTA Parallel Complement Analysis) were utilized to extract MR sequences. From the current study a total no of 210 MR sequenced were identified in Human insulin gene. The identified repeats vary in their length and majority of them are imperfect in nature. The maximum no of MR were reported in the region 3 & minimum is in the region 8 (8). In the remaining regions the no of MR sequences were lies in between maximum & minimum values. These sequences may be helpful in many molecular level studies.

## Introduction

The unique genetic makeup represents the diversity among species. This genetic pattern is denoted by arrangement of base pairs in the DNA or RNA of any species. These arrangements result in the formation of various types of repetitive sequences [1]. These sequences are found at different locations in the genomes. Their existence plays crucial roles in molecular processes happening inside the cells. The development of new methods of sequencing & other molecular technology helps to find out the different types of repeat patterns. Due to this varieties of repeats were reported in due course of time with their unique potential functions. Among them tandem repeats are one of the most common forms of repeats which shows diversity in the form of mini to microsatellite DNA. These repeats perform their role in mediating genetic plasticity as well as found to be associated with evolution [2]. VNTR is a type of tandem repeat that helps in bacterial adaptation and acts as a marker for gene mapping [3-4]. SINE & LINE sequences play their potential role in regulation of gene expression [5]. Similar to this other repeats like inverted repeats, direct repeats, indirect repeats, palindrome sequences also have unique functions in the genes or genomes [6, 7]. Some repeat patterns present at the specific site of the genomes act a signal sequence for enzymatic binding. The fundamental processes like replication, transcription & translation have involvement of such types of sequences which form the basis of their regulation [8, 9]. Many human disorders also have the involvement of repeat sequences in which their expansion causes the diseases [10, 11]. The major breakthrough in the bioinformatics area resulted in the development of tools & new pipelines to extract out the different repeat types existing in genes or genomes of the species. Such tools not only help to find out the repeat type but are also involved in studying the functional aspects of existing types [12, 13]. Among the different types of repeat sequences a unique but not much studied repeat were also identified refer to as Mirror repeat (MR). A mirror repeat always shares a center of symmetry on the same strand of the helix & forms a sequence pattern in which it looks like a mirror image [14]. Its half part is exactly similar to its rest of the portion. For example in this sequence GGATGACTTCAGTAGG, its one part GGATGACT share homology with its rest part. Mirror repeats studies shows that these sequences responsible for triple helix DNA or H-DNA formation [15]. The existence of Homopurine or Homopyrimidine mirror repeat tracks leads the formation of intramolecular triplex [16]. According to the studies it was find out that Mirror repeats also associated with diseases development [17, 18]. In the present investigation we will try to find out mirror repeats in human insulin gene sequence by utilizing a bioinformatics based approach. The current approach referred to as FPCB (FASTA PARALLEL COMPLEMENT BLAST) utilized some simple steps to extract the MR sequence from any gene or genome sequence [19]. The current study will be helpful in context of diagnosis and molecular genetics of diabetes. It will also help to study the genetic basis of variations in genes at species level.

## Methods

To identify mirror repeats, Human Insulin gene sequence (**NG_007114.1**) was analyzed using an *in silico* approach called FPCB [19]. FPCB is a manual bioinformatics based approach that helps in identification of mirror repeats. It is a three step process that involves NCBI (for gene sequence), Reverse Complement Tool (for conversion to parallel complement) and BLAST (for aligning sequence).

i. **Step 1 = Downloading gene sequence-** The sequence for the gene of interest (Human insulin gene) to be downloaded is searched on NCBI [20] and is downloaded in FASTA format. The nucleotide sequences were divided into 1000 bps regions each to make mirror identification simplified & more precise identification. ⇓
ii. **Step 2 = Making parallel complement**-After downloading the file in FASTA format and dividing it into smaller region (1000 bps) which is called our query sequence, then it would be copied into Reverse Complement Tool to obtain subject sequence (complement of query sequence). ⇓
iii. **Step 3 = Mirror repeat identification** Both the format of FASTA - query sequence and subject sequence were aligned for the BLAST homology search with some selective parameters. ⇓
iv. **Step 4 = Result and analysis-** Mirror repeat identify on the basis of position no of subject & query sequence [21].

## Results

Human insulin gene sequence were targeted under FPCB analysis to identify mirror repeats [19]. To ensure the maximum identification of Mirror repeat sequences the targeted gene sequence fragmented into 1000 bps regions. Each region denotes our original sequence which is further processed to reverse complement & BLAST analysis. A total of 210 Mirror repeat sequences were reported in the said gene sequence. The maximum no of MR sequences were reported in the region 3 (40) & minimum were in the region 8 (8). In the other regions the no of MR sequences are in between the maximum & minimum value (**Figure 1**). The complete region wise distribution of MR sequences in Human insulin gene sequence along with their length, position in the region & sequence pattern given below in the **Table 1**. The sequence patterns show that most of the sequences are imperfect in nature and varied in length also.

**Figure 1.**
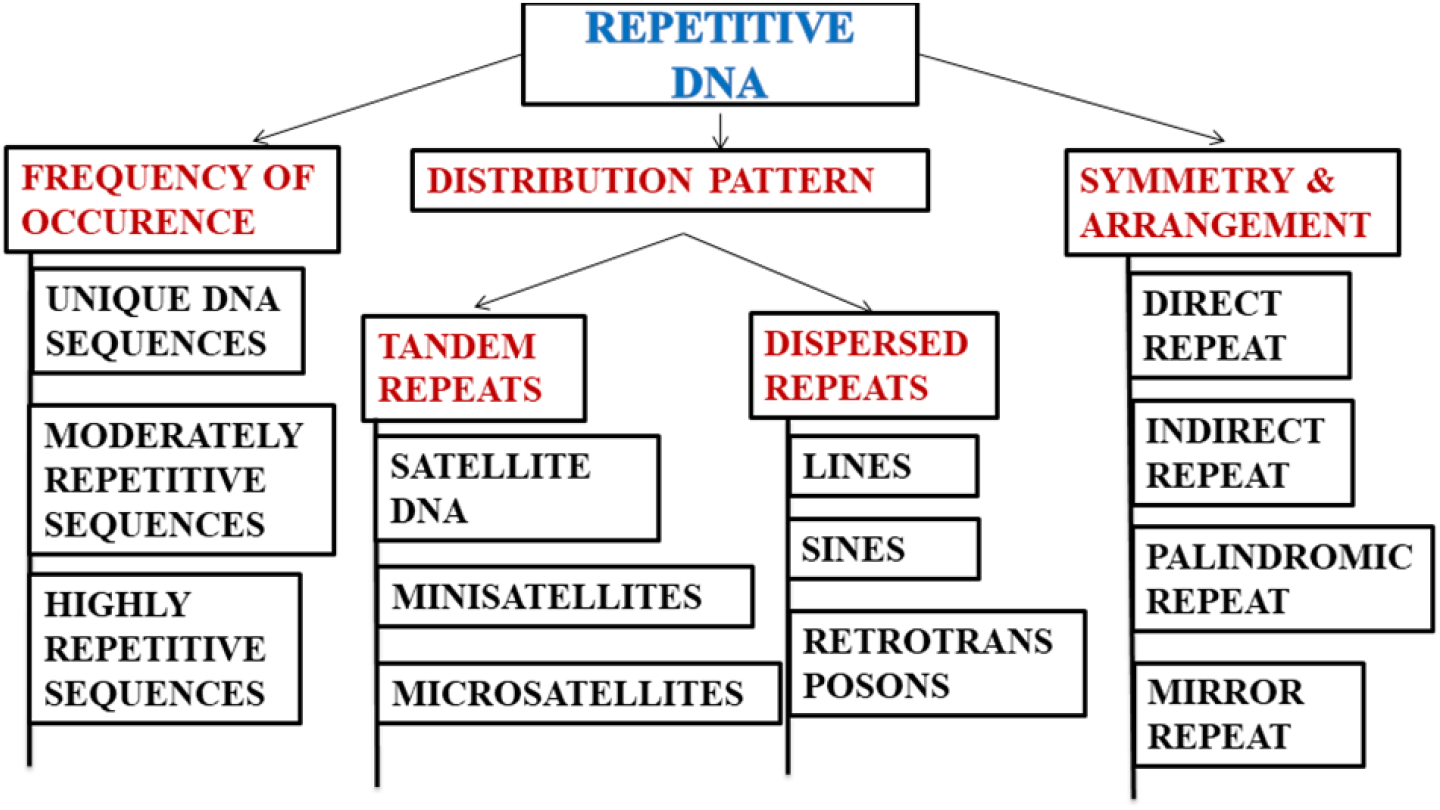
Shows various types of repeats (Adapted from **Yadav *et al* 2022 [21]**)

**Table 1-.**
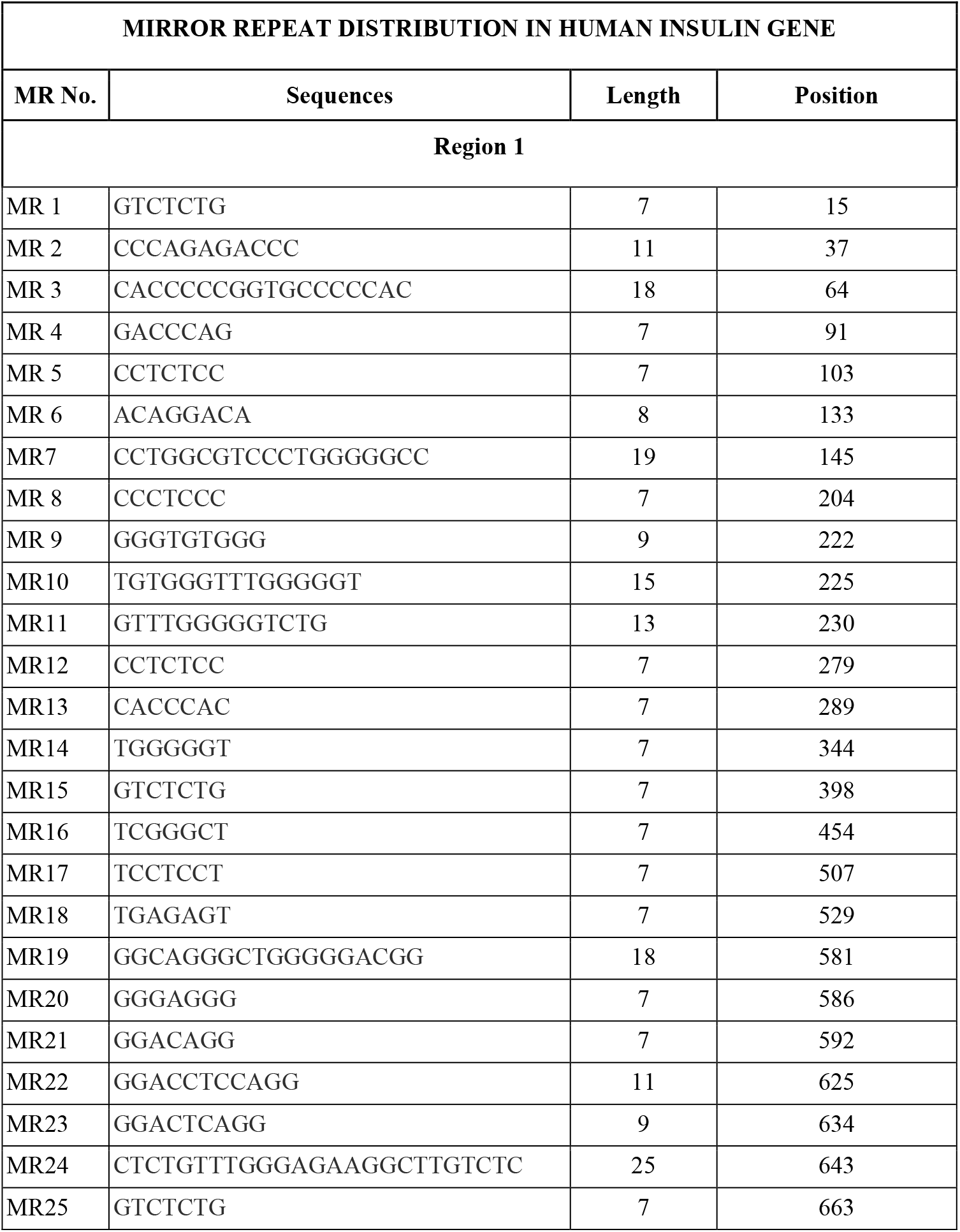
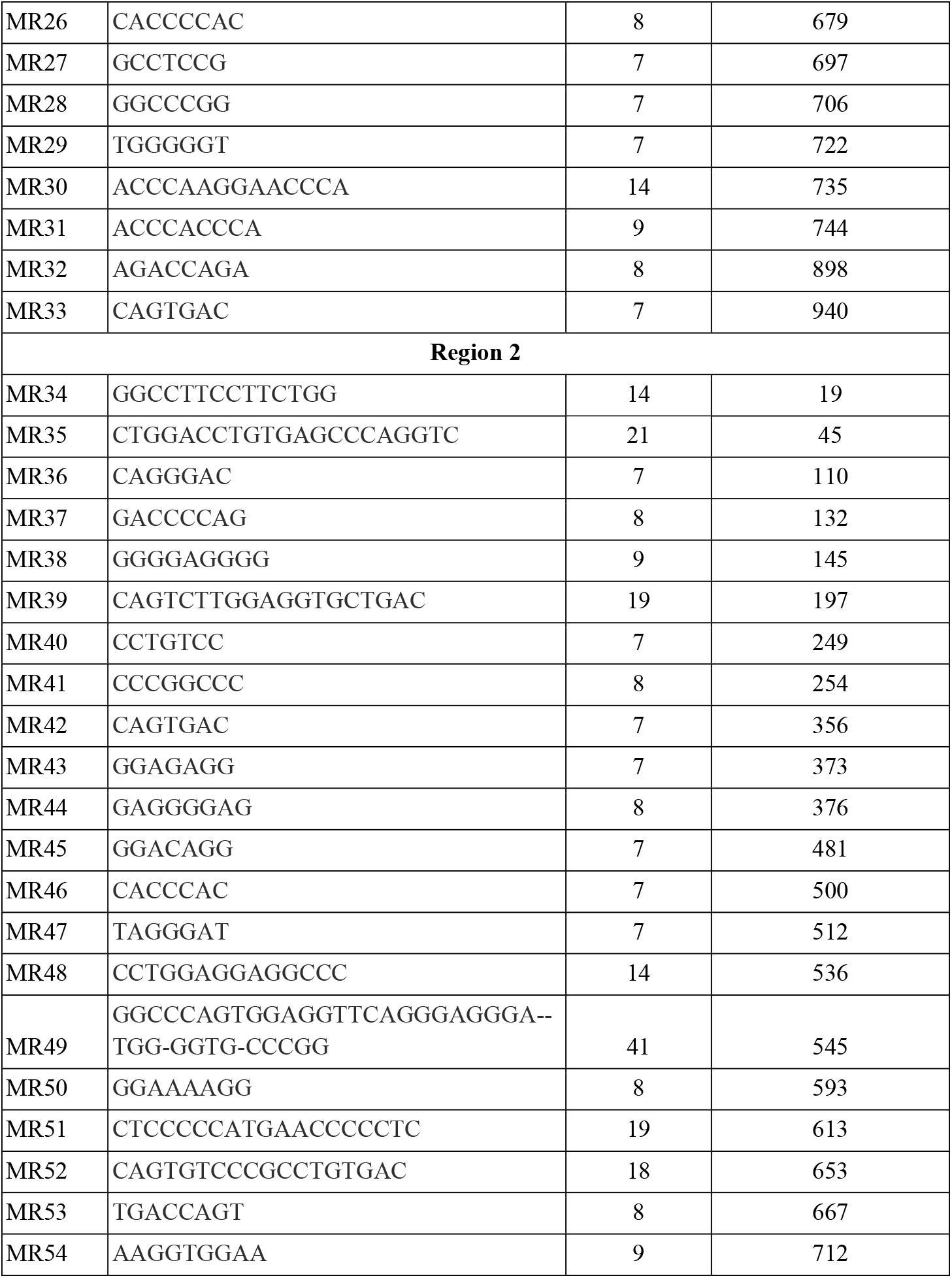
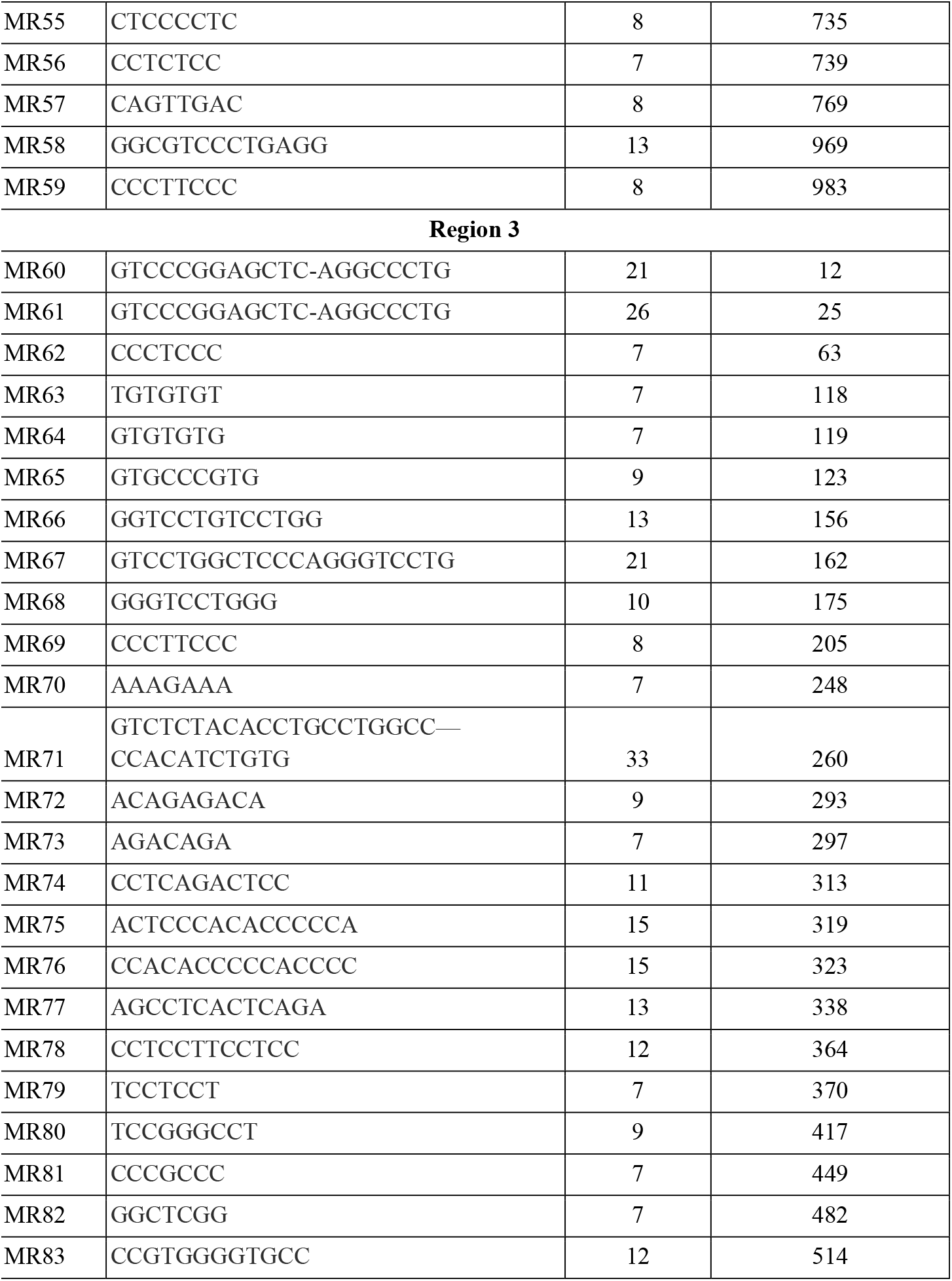
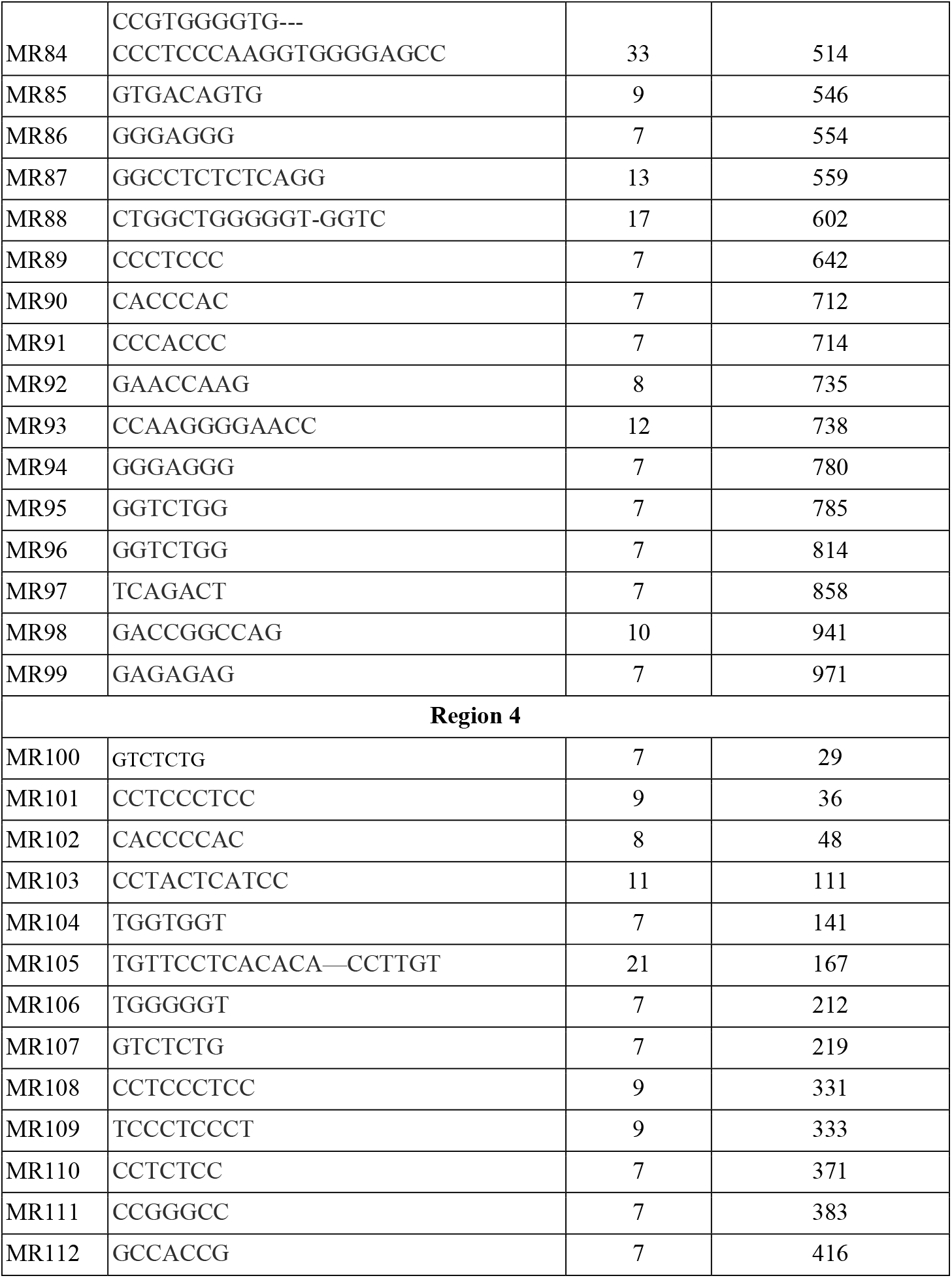
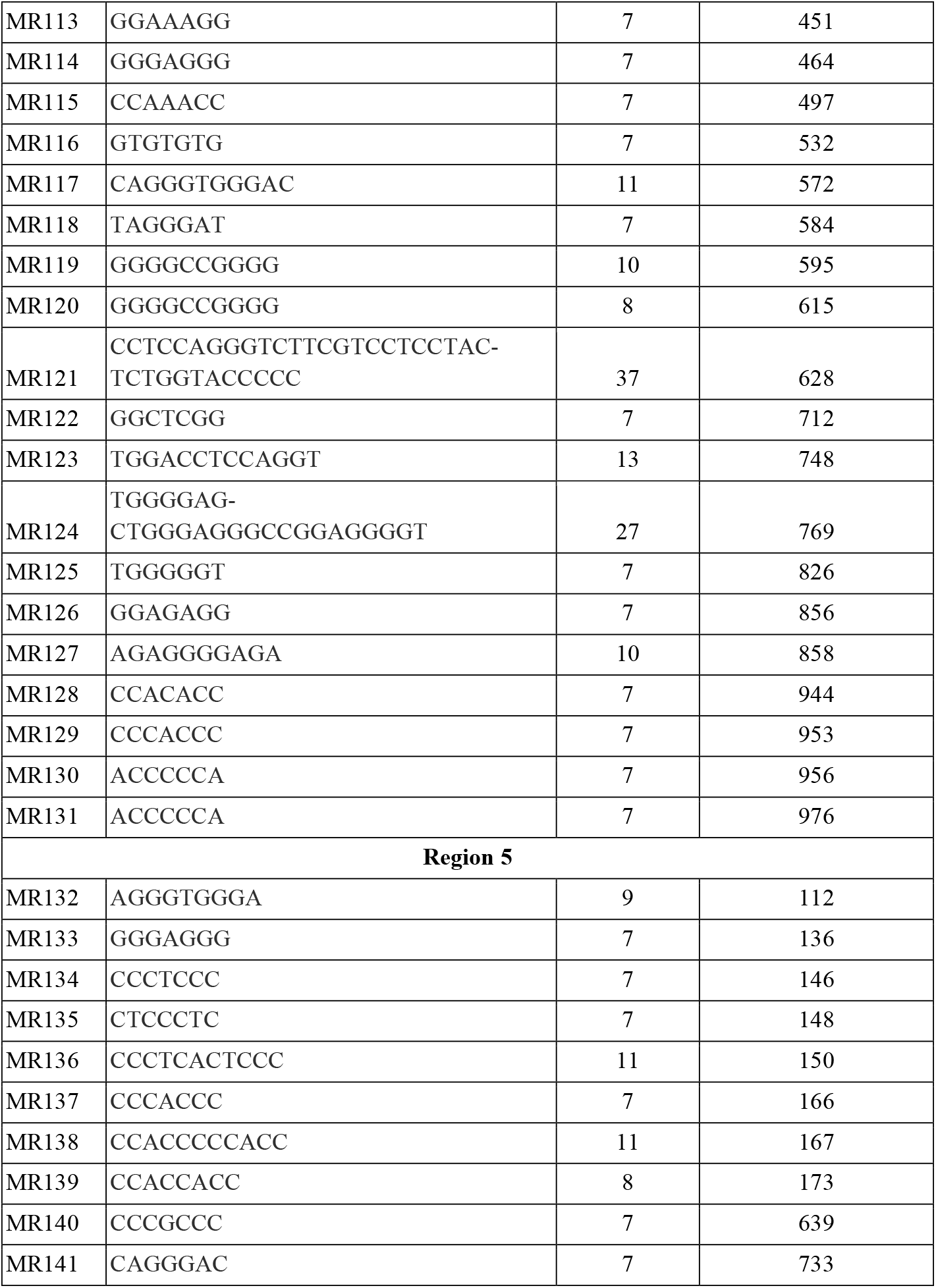
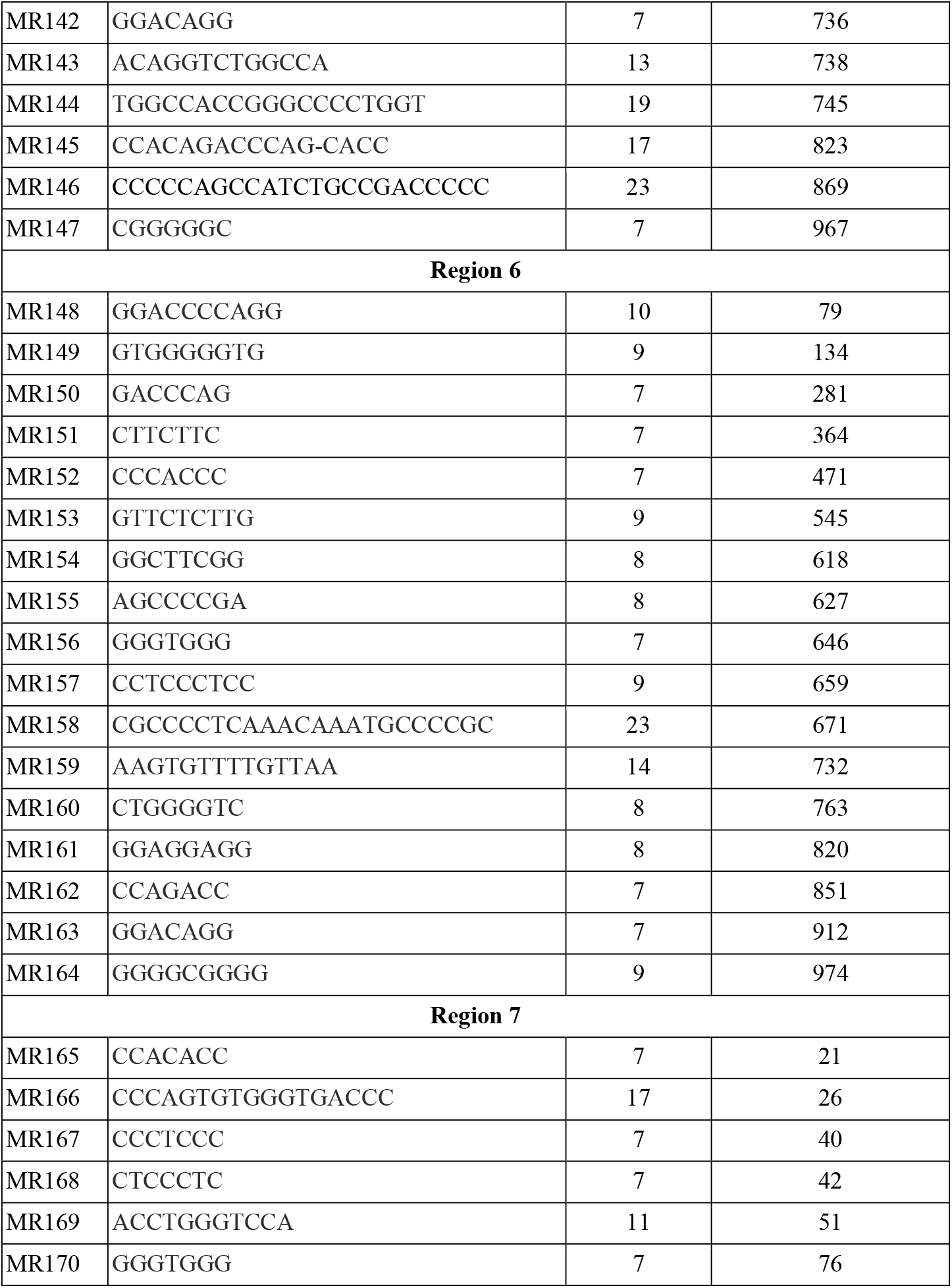
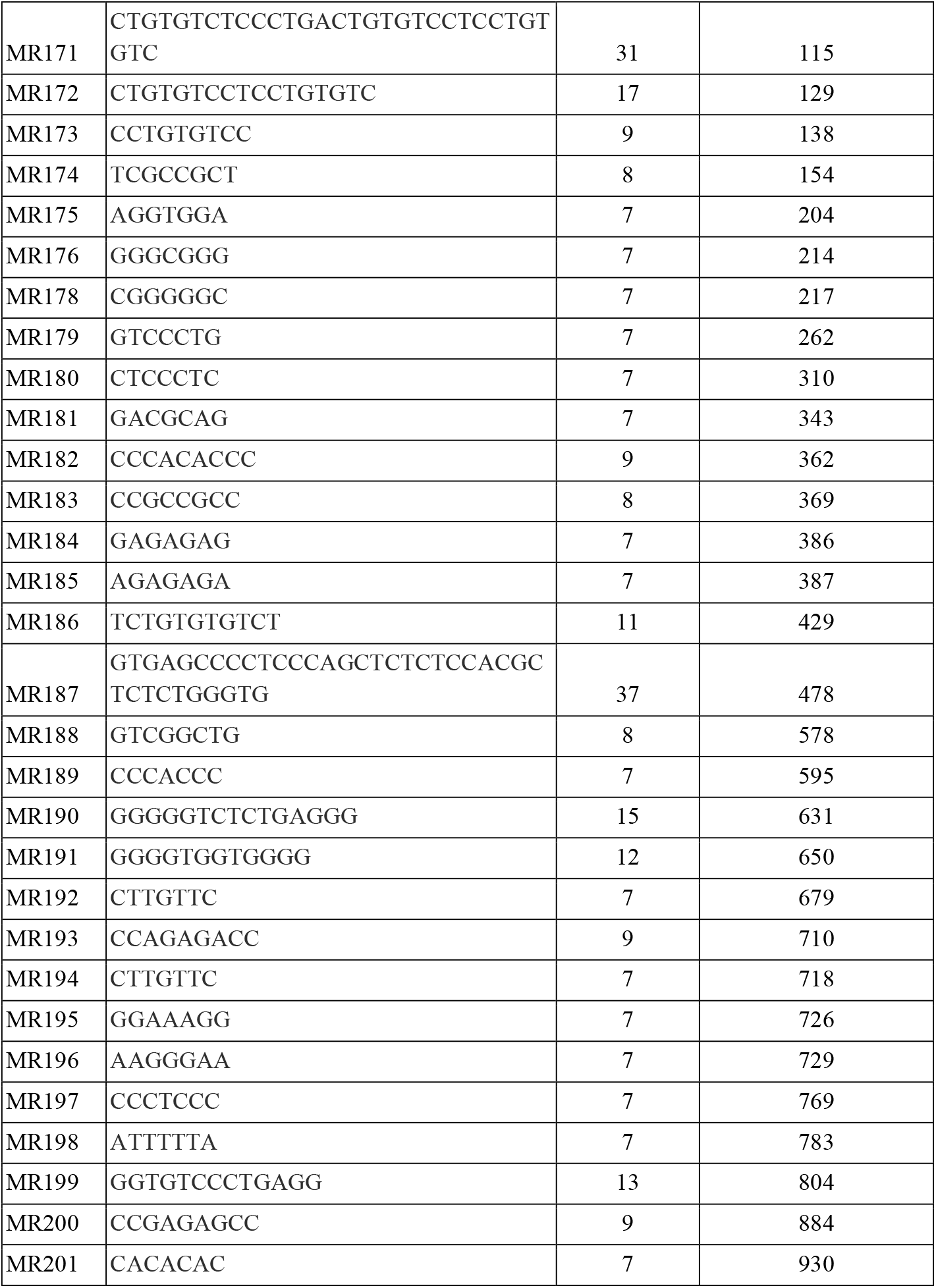
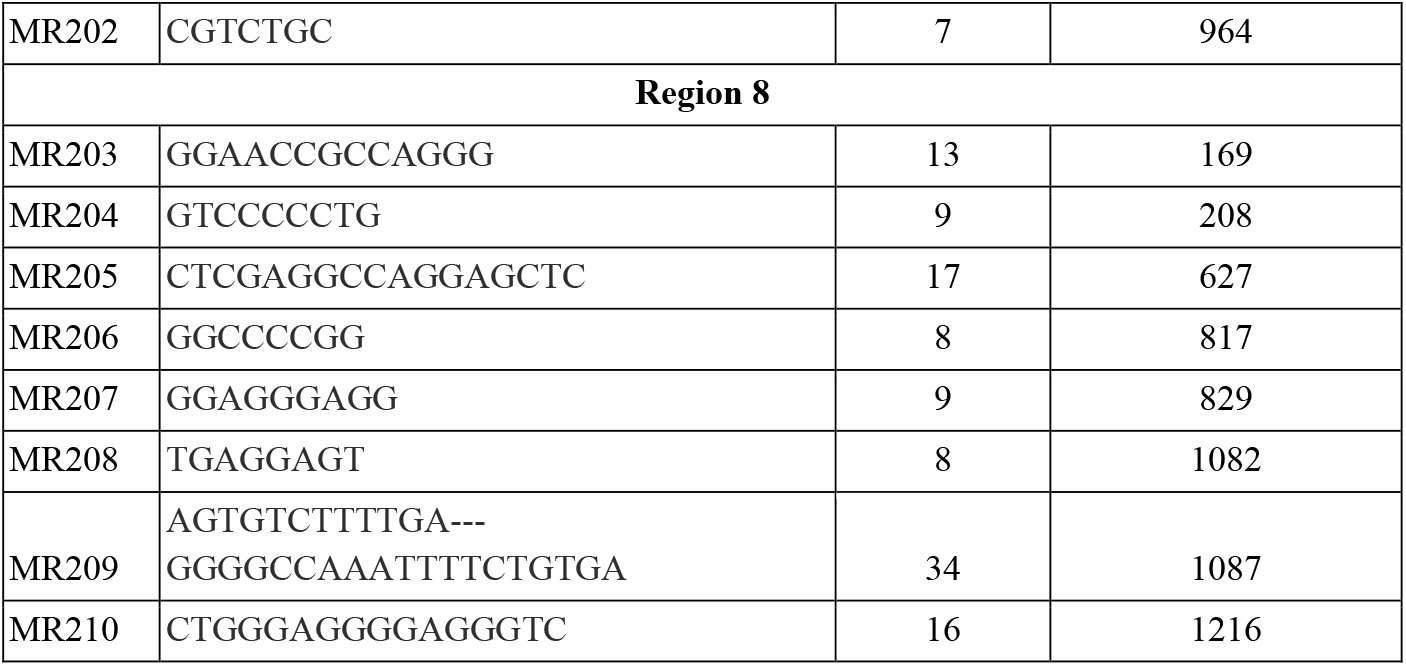
Shows Mirror repeat sequence pattern, their length & position in different regions of Human Insulin gene sequence

**Figure 2.**
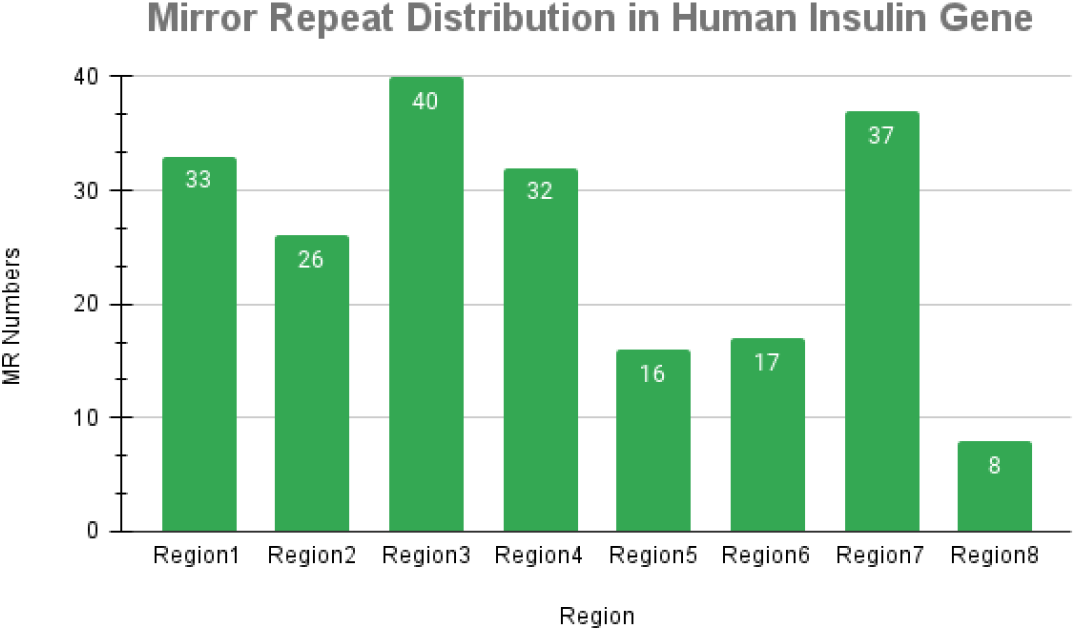
Shows Mirror repeat distribution in Human Insulin Gene sequence. The maximum & minimum no of MR sequence were reported in **Region 3 (40) & Region 8** (8) respectively.

## Discussion

The present study was also supported by previous ones in which Mirror repeats patterns were observed in case of HIV-1 & HIV-2 genome. A total no of 232 & 248 MR sequences were identified in case of HIV using FPCB analysis [22]. The similar studies carried out using *E*.*coli* genes [23], model plant [21, 24] as well as viral domains [25] shows occurrence of MR sequences throughout the genome or gene sequences. According to the present study their existence in the human insulin gene was also an indicator that these sequences may be has some significant role in the targeted gene. Some *in silico* methods can be utilized in biological sciences to check their significance in relation to clinical prospective [26, 27]. In tools & techniques based study these sequences will be utilized as a biological marker for testing. These can also compare with respect to individual/species & may be treated as an indicator marker to check the molecular evolution of the different domains. This will be a novel way to classify the species if the conservative MR sequences are existed in the gene or genome sequences. In modern emerging areas of bioinformatics this study will be helpful in context of finding & designing the new tools for MR study.

## Conclusion

Bioinformatics based analysis concluded here that the occurrence of mirror repeats in Human Insulin Gene sequence will provide a hint about their key roles at molecular level. Their distribution throughout the gene can be an indicator of their multiple uses which will be in study of diseases development, evolution & mutation bases study, diagnosis & therapeutic based studies etc.

## Acknowledgment

The authors of this manuscript want to acknowledge managing authority of Starex University to provide their sincere support to carry out this study.

## Author contribution statement

The whole experiments & analysis carried out by Ms. Manisha Yadav & Ms. Barkha Sehrawat, both authors contribute equally. Ms. Pooja Yadav, Ms. Priyanka Yadav, Ms. Jyoti Kumari, Ms. Rachna Yadav, Ms. Amrita Singh, Ms. Shivani Yadav help Ms. Manisha Yadav & Barkha Sehrawat in data analysis as well as in the writing & drafting of the manuscript. Dr. Sandeep Yadav (Corresponding author) help in revision & finalized the final version of this manuscript. All authors approved the final version of this manuscript.

## Conflict of Interest

None

## Funding Acknowledgment

No any financial support received from any funding agency for this work.

## Ethical Statement

No any method used unethically in the present work.

## Notes

### Competing Interest Statement

The authors have declared no competing interest.

